# RecO impedes RecG-SSB binding to impair the strand annealing recombination pathway in *E.coli*

**DOI:** 10.1101/708271

**Authors:** Xuefeng Pan, Li Yang, Nan Jiang, Xifang Chen, Bo Li, Xinsheng Yan, Yu Dou, Liang Ding, Fei Duan

## Abstract

Faithful duplication of genomic DNA relies not only on the fidelity of DNA replication itself, but also on fully functional DNA repair and homologous recombination machinery. We report a molecular mechanism responsible for deciding homologous recombinational repair pathways during replication dictated by binding of RecO and RecG to SSB in *E.coli.* Using a RecG-yfp fusion protein, we found that RecG-yfp foci appeared only in the Δ*rec*G, Δ*rec*O and Δ*rec*A, Δ*rec*O double mutants. Surprisingly, foci were not observed in wild-type Δ*rec*G, or double mutants where *recG* and either *recF* or, separately *recR* were deleted. In addition, formation of RecG-yfp foci in the Δ*rec*O::kan^R^ required wildtype *ssb*, as *ssb-113* could not substitute. This suggests that RecG and RecO binding to SSB is competitive. We also found that the UV resistance of *rec*O alone mutant increased to certain extent by supplementing RecG. In an *ssb-113* mutant, RecO and RecG worked following a different pattern. Both RecO and RecG were able to participate in repairing UV damages when grown at permissive temperature, while they could also be involved in making DNA double strand breaks when grown at nonpermissive temperature. So, our results suggested that differential binding of RecG and RecO to SSB in a DNA replication fork in *Escherichia coli*.may be involved in determining whether the SDSA or DSBR pathway of homologous recombinational repair is used.

**Author summary:** Single strand DNA binding proteins (SSB) stabilize DNA holoenzyme and prevent single strand DNA from folding into non-B DNA structures in a DNA replication fork. It has also been revealed that SSB can also act as a platform for some proteins working in DNA repair and recombination to access DNA molecules when DNA replication fork needs to be reestablished. In *Escherichia coli*, several proteins working primarily in DNA repair and recombination were found to participate in DNA replication fork resumption by physically interacting with SSB, including RecO and RecG etc. However the hierarchy of these proteins interacting with SSB in *Escherichia coli* has not been well defined. In this study, we demonstrated a differential binding of RecO and RecG to SSB in DNA replication was used to establish a RecO-dependent pathway of replication fork repair by abolishing a RecG-dependent replication fork repair. We also show that, RecG and RecO could randomly participate in DNA replication repair in the absence of a functional SSB, which may be responsible for the generation of DNA double strand breaks in an *ssb*-113 mutant in *Escherichia coli.*

## Introduction

Genome duplication is highly processive and accurate, relying not only on DNA replication itself, but also on the proper cooperation between DNA replication, repair and homologous recombination. This is particularly true when DNA replication encounters DNA lesions, including damaged bases, single and/or double-strand breaks, DNA strand templates bound by proteins etc. [1-10]. In the last decades, the roles of certain homologous recombination proteins including RecG, RecQ, RuvAB, PriA, PriB, RecA and RecFOR have become increasingly clear [3, 11-17]. In addition, it has also become clear that the single-strand binding protein (SSB) plays critical roles in repair processes as it both binds to unwound strands of the duplex affording protection and also binds to as many as fourteen proteins that comprise the SSB interactome (Lecoite and Shereda papers as reference). Included in the partner list are the DNA helicase RecG and the RecA-mediator, RecO, both of which are the focus of this study (refs: YU for RecG and Korolev & Bianco for RecO).

In *E.coli*, RecG is a 76 kDa monomeric protein that has both double-stranded DNA helicase/translocase activity and RNA: DNA helicase activity. RecG has been found to take part in multiple pathways of DNA metabolism, including homologous recombination pathways such as RecBCD, RecF and RecE pathways [12, 13, 18, 19]; regression of stalled DNA replication fork under UV irradiations by annealing the two nascent strands in the leading and lagging template, respectively [14, 16-24]; avoidance of stable RNA-DNA hybrid in transcription [20, 21]; conversion of 3-stranded junctions into 4-stranded ones [12,25,26]; catalyzing migration of Holliday junctions to facilitate their resolution [3,20]; promoting and opposing RecA-catalyzed strand exchange [3,27]; processing DNA flaps made when DNA replication forks converge[28] and stabilizing D-loops[20]. The roles of RecG in *E.coli* cells have been reconciled as to assist PriA positioning correctly in a D-loop structure to re-establish a DNA replication fork [11, 29]. Interestingly, a storage form of RecG, made up of RecG, PriA and SSB has recently been reported in *E.coli* cells, whose formation requires that SSB was present in excess over the RecG and PriA, and also intact C-terminus of SSB, but does not need the presence of any DNA molecules [30].

RecG comprises three functional domains, amongst which, domain I (wedge domain) which binds nucleic acid, and domains II and III which comprise the helicase domains and bind both duplex DNA and ATP [13, 31, and 32]. RecG is targeted to replication fork-like structures via interacting with the C-terminus of SSB, forming a stoichiometric RecG-SSB complex by 2 RecG monomers binding to an SSB tetramer [32]. SSB-113 whose proline residue of 176 in the C-terminus of the SSB was substituted by serine, binds poorly to RecG [33, 34]. The binding of SSB can transiently inhibit the helicase activity of RecG, while assisting the translocation of RecG to the parental double stranded DNA region of the DNA replication fork substrate [32].

RecO is a 27 kDa protein that binds single-and double-stranded DNA via its oligonucleotide-binding fold (OB-fold) in the N-terminal domain [35]. It catalyzes single stranded DNA annealing of complementary oligonucleotides complexed with SSB in an ATP-independent manner [36-39]. RecR can inhibit the annealing activity of RecO by binding to it, forming RecOR complex [22, 34, 40-42]. In homologous recombination, RecOR facilitates the initiation and growth of RecA-ssDNA by forming a RecO-SSB-ssDNA intermediate, and by RecR inhibiting the strand annealing activity of RecO, while activating the ssDNA-dependent ATPase activity of RecA [3, 35, 38, 39, 43-54]. In addition, the concerted actions of RecO, RecR and RecF were found to protect a stalled DNA replication fork from degradation of the lagging strand template by RecQ and RecJ under UV irradiation [49, 50, 53, and 54].

So far, more than 14 proteins, including the X subunit in DNA polymerase III holoenzyme, DNA polymerase V, RecQ, PriA, PriB, RecG, RecO, ExoI etc. have found to be SSB interacting proteins [55-64]. Some of them interact with the unstructured C-terminus of SSB when working at the interface of DNA replication and recombination [34, 42, 62, and 65]. The *E.coli* SSB forms a homotetrameric protein complex that plays a central role in DNA replication, repair and recombination [60, 61, and 65]. In DNA replication, SSB binds single stranded DNA (ssDNA) of the lagging strand template in a DNA replication fork with no DNA sequence specificity, protecting the ssDNA from misfolding and also from nuclease attack[67,68]. In homologous recombination, SSB prevents recombinase RecA from loading onto ssDNA until recombination mediator proteins (RMPs) come to facilitate the exchange of RecA with SSB for ssDNA.

Recently SSB proteins isolated from both *E.coli* and *B.subtilis* have found to bear unstructured C-terminal tail domains (SSB-Cter) that are responsible for the interacting with the aforementioned proteins. In *E.co*li, deletion of the last 8 amino acids of the SSB-Cter made cells unviable, and even a point mutation e.g. *ssb-113* greatly compromises the cell growth, likely by impairing interactions with interactome partners. Importantly, proteins interacting with the SSB-Cter can gain access to the DNA replication fork as shown recently for RecG [32].

In this work we have attempted to determine how RecG and RecO are utilized in the presence or absence of SSB binding in live *E.coli* cells. To this end, we visualized the RecG protein fluorescent focus formation in the cells carrying RecO, RecR or RecF null mutant genes, respectively. We also compared the growth and DNA repair capacities of the mutants. We found for the first time that binding of RecG to SSB-Cter in wildtype *E.coli* is impaired by RecO, possibly by direct competition. These results suggest that RecO and RecG can work differently in the growth and UV damage repair depending on SSB-binding.

## Results

### Experimental Rationale

RecG and RecO bind to the unstructured C-terminus of SSB *in vitro* [33, 46]. However, visualization of RecG and RecO molecules in a DNA replication fork in *vivo* has been unsuccessful so far [64, 69]. We hypothesized that a competition between RecG and RecO binding to SSB-ssDNA via the C-terminus of SSB could play a critical role in regulating pathways of DNA metabolisms at the interface of DNA replication, repair and homologous recombination when DNA replication turned out to be difficulty due to some situations aforementioned.

To test this possibility, we constructed the RecG and RecG-yfp expression plasmids, *rec*G-pUC18 and *recG-yfp*-pUC18 (Figure 1A and 1B). In parallel a set of *E.coli* isogenic mutants such as JM83 Δ*rec*G265::*Cm*^*R*^ and corresponding *recF, O and R* deletion mutants were constructed by P1 transduction. Next, we verified that the RecG and RecG-yfp were constructed correctly by DNA sequencing and found to be functional by determining their ability to complement the Δ*rec*G265::*Cm*^*R*^ mutant using UV irradiation. We found that the UV resistance of the JM83Δ*rec*G265::Cm^R^/*rec*G-pUC18 and separately, JM83Δ*rec*G265::*Cm*^*R*^/*recG-yfp*-pUC18 were improved, albeit only partially (Figure 1C), showing that both RecG and RecG-yfp expressed by *rec*G-pUC18 and *recG-yfp*-pUC18 plasmids were functional.

**Figure 1.**
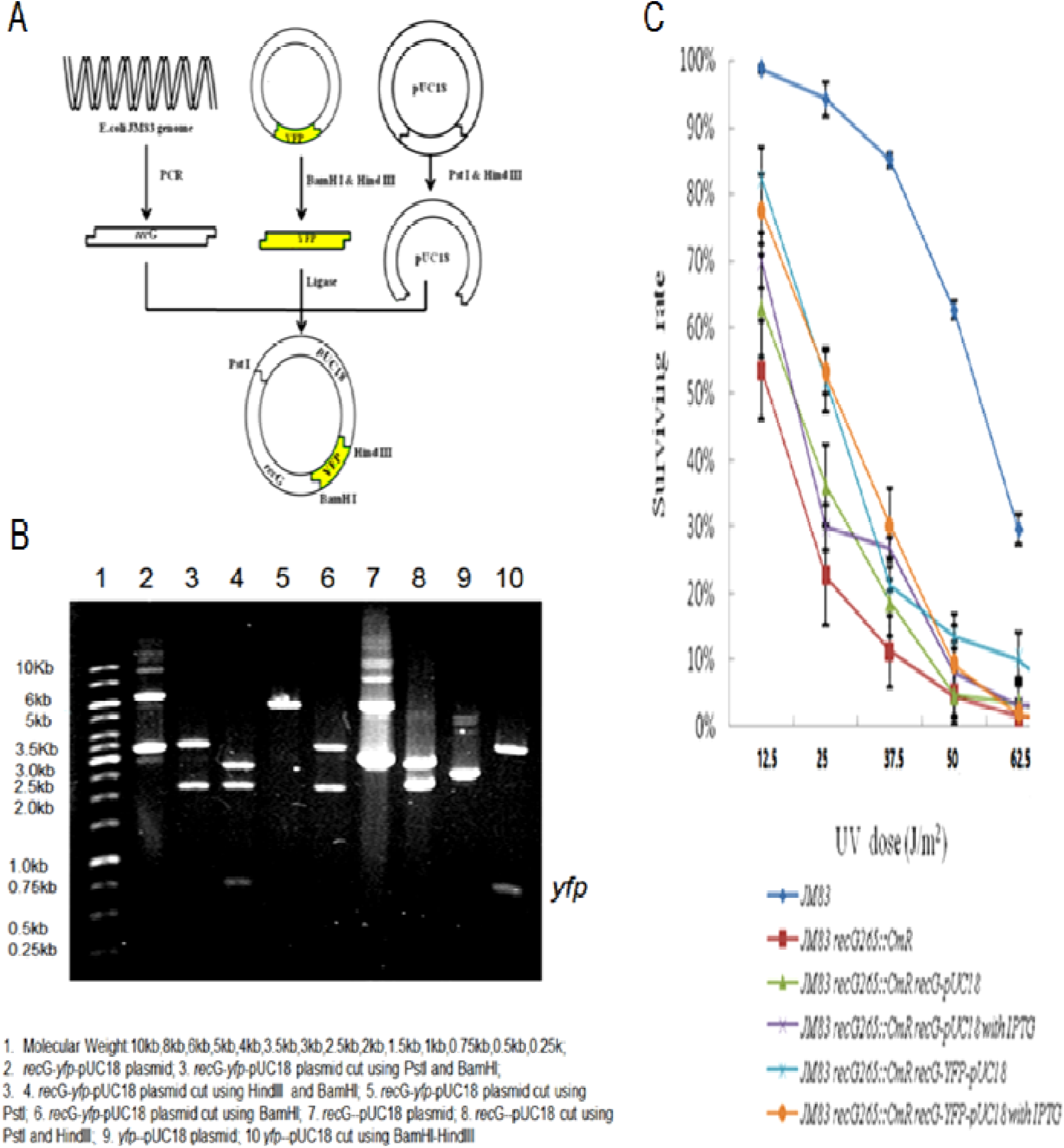
Construction of plasmids. 1A, schematic illustration for *RecG-yfp*-pUC18 plasmid construction 1B, restriction mapping analysis of *RecG-yfp*-pUC18 and *rec*G--pUC18 plasmids 1, molecular Weight: 10kb, 8kb, 6kb, 5kb, 4kb, 3.5kb, 3kb, 2.5kb, 2kb, 1.5kb, 1kb, 0.75kb, 0.5kb, 0.25k; 2, *RecG-yfp*-pUC18 plasmid; 3. *RecG-yfp*-pUC18 plasmid cut using *Pst*I and *BamH*I; 4, *RecG-yfp*-pUC18 plasmid cut using *Hind*III and *Bam*HI; 5. *RecG-yfp*-pUC18 plasmid cut using *Pst*I; 6. *RecG-yfp*-pUC18 plasmid cut using *Bam*HI; 7. *rec*G-pUC18 plasmid; 8. *rec*G-pUC18 cut using *Pst*I and *Hind*III; 9. *yfp*-pUC18 plasmid; 10 *yfp*-pUC18 cut using *Bam*HI-*Hind*III C, comparison on UV resistance of JM83 strains carrying either *RecG-yfp*-pUC18 or *rec*G--pUC18 plasmid

### RecG-yfp did not form fluorescent foci in the wild-type and the Δ*recG265::Cm*^*R*^ mutant

We then visualized the RecG-yfp protein in the wildtype JM83 and the JM83Δ*rec*G265::Cm^R^ mutant using confocal microscopy. This was done by transformation with *RecG-yfp*-pUC18 and *yfp*-pUC18, respectively (Figure 2). We found that in all cases, the YFP signal was found scattered throughout the cytoplasm with no foci being readily observed (Figure 2).

**Figure 2.**
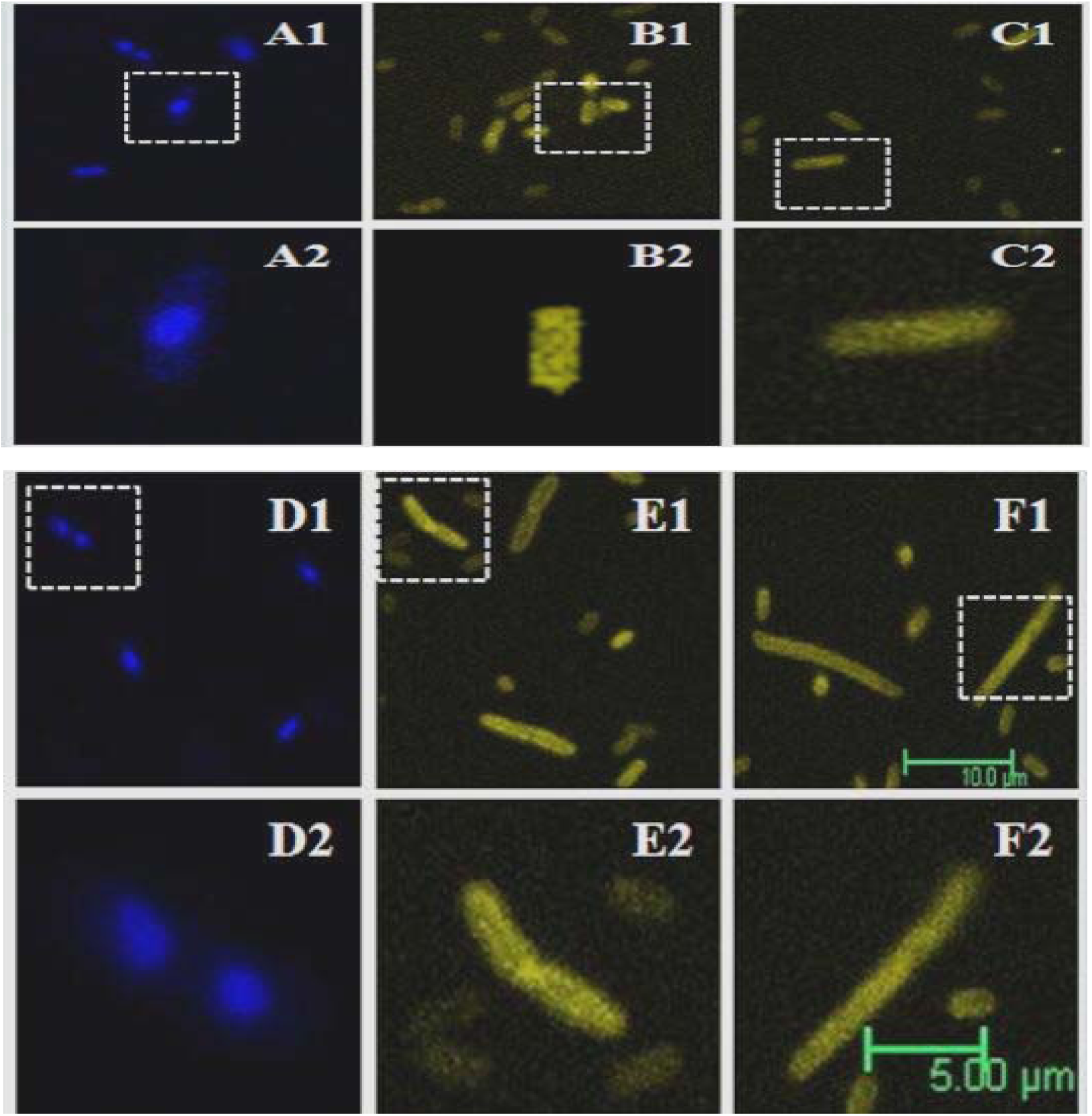
Visualization of the RecG-yfp in a JM83 *rec*G::Cm^R^ mutant. A1,A2: JM83 stained using DAPI; B1,B2:JM83(*RecG-yfp*-pUC18); C1,C2:JM83(*yfp*-pUC18); D1,D2: JM83Δ*rec*G265::*Cm*^*R*^ stained using DAPI; E1,E2: JM83Δ*rec*G265::Cm^R^(*RecG-yfp*-pUC18); F1,F2: JM83Δ*rec*G265::Cm^R^(*yfp*-pUC18)

### RecG-yfp formed fluorescent foci only when RecO was absent

The inability to form RecG-yfp foci in the JM83Δ*rec*G265::Cm^R^ was surprising in light of previous results (Lloyd paper in NAR – RecG localizes to forks). We surmised that this might be due to the presence of one or more inhibitors. To identify the inhibition factor(s), we then visualized RecG-yfp protein in isogenic strains where (i) both *rec*G and *rec*F were deleted; or separately, (ii) *rec*G and *rec*O and finally (iii), *rec*G and *rec*R. Surprisingly, we found that RecG-yfp foci could be visualized only in the JM83Δ*rec*G265:: CmRrecO::Kan^R^ mutant cells. In fact, foci were evident in 50∼60% of the cells (Figure 3A, I1 and I2, Figure 3B). In contrast, foci were not observed in the *rec*G/*rec*F and *rec*G/*rec*R double mutants (Figure 3G1, G2, K1, and K2).

**Figure 3.**
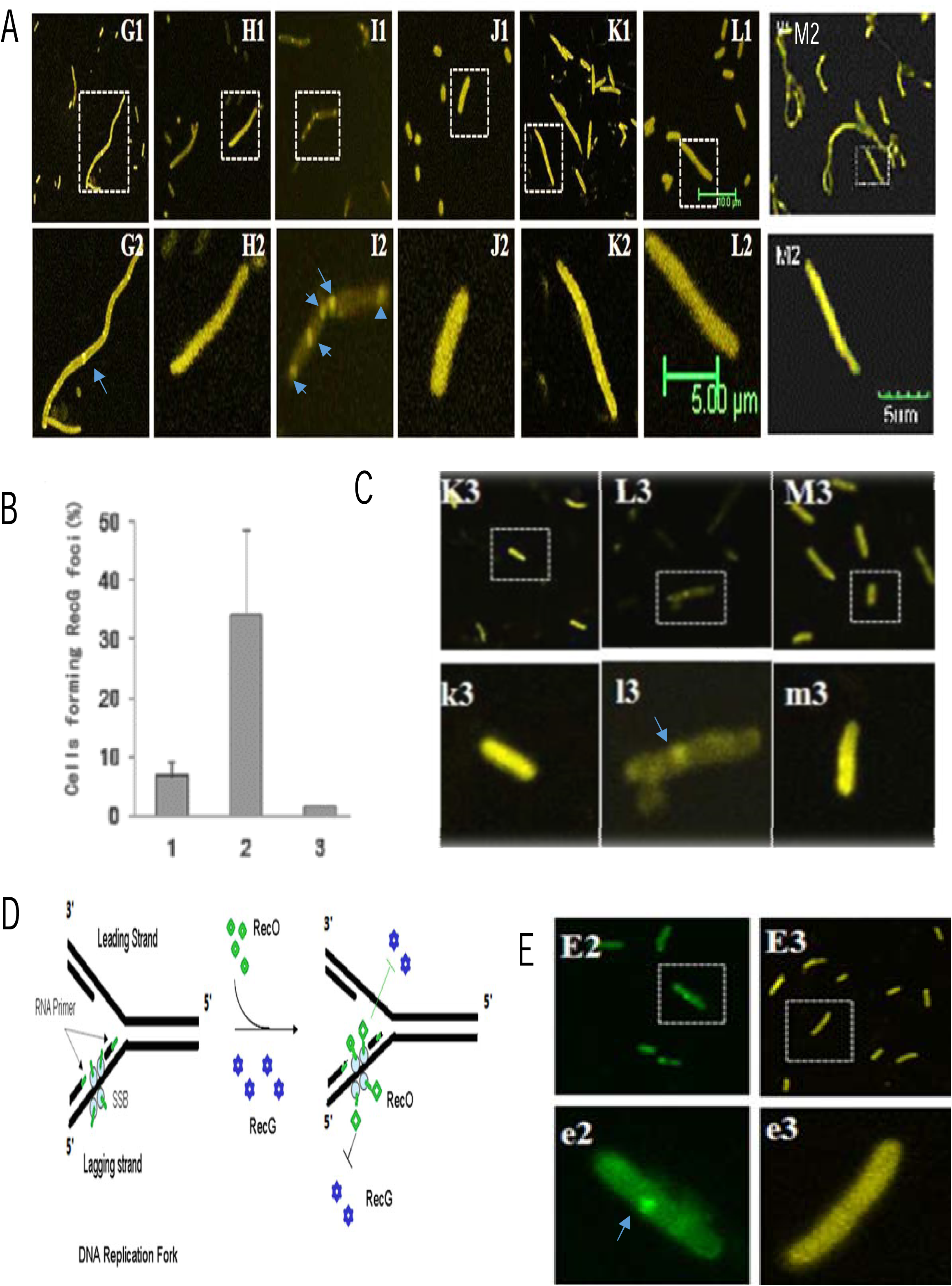
Visualizations of RecG-yfp protein in *rec*F, *rec*R, *rec*O, *ssb-113* mutants. *3A*, visualizations of RecG-yfp proteins in the JM83 strains with the absence of RecF, RecR, RecO and the presence of SSB-113 G1,G2: JM83*rec*F::Kan^R^Δ*rec*G265::Cm^R^ (*RecG-yfp*-pUC18); H1,H2:JM83*rec*F::Kan^R^Δ*rec*G265::Cm^R^(*yfp*-pUC18); I1,I2:JM83*rec*O::Kan^R^Δ*rec*G265::Cm^R^(*RecG-yfp*-pUC18); J1,J2:JM83*rec*O::Kan^R^Δ*rec*G265::Cm^R^(*yfp*-pUC18); K1,K2:JM83*rec*R::Kan^R^Δ*rec*G265::Cm^R^ (*RecG-yfp*-pUC18); L1,L2: JM83*rec*R::Kan^R^Δ*rec*G265*::Cm*^*R*^(*yfp*-pUC18) M1,M2: JM83*rec*O::Kan^R^Δ*rec*G265::Cm^R^*ssb-113*(ts)(*RecG-yfp*-pUC18) **3B, the ratio of the RecG-yfp foci in the nucleoid** 1, JM83*rec*F:: Kan^R^Δ*rec*G265::*Cm*^*R*^ (*recG-yfp*-pUC18) 2, JM83*rec*O:: Kan^R^Δ*rec*G265::Cm^R^ (*recG-yfp*-pUC18) 3, JM83*rec*R:: Kan^R^ Δ*rec*G265::Cm^R^ (*recG-yfp*-pUC18) **3C, Visualizations of RecG-yfp fusion proteins in double mutants of JM83recF:: Kan**^**R**^ Δ ***rec*A::Cm**^**R**^, **JM83*rec*O:: Kan**^**R**^ Δ***rec*A::Cm**^**R**^ **and JM83*rec*R:: Kan**^**R**^ Δ***rec*A::Cm**^**R**^ K3, k3: JM83*rec*F:: Kan^R^Δ*rec*A::Cm^R^ (*recG-yfp*-pUC18) L3, l3: JM83*rec*O:: Kan^R^Δ*rec*A::Cm^R^ (*recG-yfp*-pUC18) M3, m3: JM83*rec*R:: Kan^R^ Δ*rec*A::Cm^R^ (*recG-yfp*-pUC18) **3D, visualizations of MutS-gfp and RecR-yfp protein focus in a JM83*mut*S::Tc**^**R**^***rec*O::Kan**^**R**^ **double mutant** E2, e2: JM83*mut*S::Tc^R^*rec*O::Kan^R^(*mutS-gfp*-pACYC184); E3, e3: JM83*mut*S::Tc^R^*rec*O::Kan^R^(*recR-yfp*-pUC18)

To determine whether the foci were due to RecG-yfp protein aggregates, we also visualized their distribution in a set of isogenic double mutant strains: Δ*rec*F/Δ*rec*A, Δ*rec*O/Δ*rec*A and Δ*rec*R/Δ*rec*A. These strains are sensitive to UV irradiation, showing reduced viability but are reduced for filamentation. As before, foci were only observed in the Δ*rec*O background, but their number was reduced compared to the Δ*rec*O/Δ*rec*G strain (Figure 3C).

We interpreted this decrease in focus formation in JM83 Δ*rec*O::Kan^R^Δ*rec*A::Cm^R^ to two possible reasons. First, filamentous growth of JM83Δ*rec*G265::Cm^R^ *rec*O::Kan^R^ (as shown in Figure 3A) was prevented by deactivating *rec*A, preventing multiple nucleoid accumulation in the cell. Second, there may be competition between the wildtype and fusion RecG in the *rec*A/*rec*O double mutant, even though RecG presents at low levels (less than 10 molecules per cell) [31].

Our results showing RecG-yfp focus formation in only the Δ*rec*G265::Cm^R^ *rec*O::Kan^R^ and ΔrecO::Kan^R^Δ*rec*A::Cm^R^ strains suggested their inhibition in *recO*^*+*^ cells is due to RecO protein rather than by RecF and RecR (Figure 3A, C and D).

In addition, we also carried out a validating visualization analysis on the distributions of RecR-yfp and MutS-gfp in the cells carrying both *rec*O and *mut*S mutations. This makes sense as MutS binds both the mismatched base pair in double stranded DNA and the beta clamp in a DNA polymerase III holoenzyme, and RecR-yfp appears in the nucleoid region in *E.coli* [70]. We expressed the MutS-gfp by using *MutS-gfp*-pACYC184 and RecR-yfp by *rec*R-*ypf*-pUC18 plasmids in a JM83 *rec*O::Kan^R^*mut*S::Tc^R^ mutant. These results show that MutS-gfp focus formation was similar to that of RecG-yfp in the Δ*rec*O/Δ*rec*A (Fig 3E). In contrast, RecR-yfp did not form detectable foci, consistent with the finding that *E.co*li RecR alone does not bind DNA and RecR-yfp formed foci through indirect binding to nucleoid as shown previously [70]. Although we noticed that RecG and PriA also formed protein complex with SSB near the inner membrane in *E.co*li cells, which did not require DNA molecules [30], we visualized the RecG-yfp fluorescent foci in the strains of JM83Δ*rec*G265::Cm^R^ *rec*O::Kan^R^ and JM83Δ*rec*O::Kan^R^Δ*rec*A::Cm^R^ where the *pri*A gene remains unaffected. Therefore, we understood that our observation may suggest a possible hindrance of RecG by RecO in binding to SSB at the interface of DNA replication, repair and homologous recombination involving RecO and RecG.

### Forming RecG-yfp foci in the RecO mutant required a functional SSB C-terminus

To further understand if the RecG-yfp protein foci formed in JM83Δ*rec*G265::Cm^R^ *rec*O mutant required RecG interacting with the C-terminus of SSB, we then constructed a JM83Δ*rec*G265::Cm^R^ *rec*O*ssb*-*113*^(ts)^ triple mutant (an *E.coli* mutant with complete deletion of C-terminus of SSB was unviable). The *ssb*-113 gene encodes a SSB-113 mutant protein carrying a P176S substitution at the C-terminus of SSB [71]. Such an SSB-113 mutant retained ssDNA binding capacity and allows DNA replication to occur albeit at lower temperature. Importantly, it shows diminished multiple protein interactions with its unstructured C-terminus, including X subunit of DNA polymerase III, PriA, RecQ, ExoI, PolV, RecG, RecJ, RecO, sbcC and Ung etc. [33, 34, 55, 57, 58, 59, 60,62, 72], making such mutants to be sensitive to higher temperature, UV irradiation because of generation of DNA double strand breaks [33, 34, 55,57, 58, 59, 60, 62, 67].

We then visualized the RecG-yfp fluorescent focus formation by transforming *rec*G-*yfp*-pUC18 plasmid into the Δ*rec*G*rec*O*ssb*-113^(ts)^ triple mutant. Once again, we found that the RecG-yfp proteins expressed in the cytoplasm of JM83Δ*rec*G265::Cm^R^ *rec*O*ssb*-113^(ts)^ at permissive temperature (30°C) were scattered (Figure 3A, M1 and M2), when compared with those RecG-yfp proteins expressed in JM83Δ*rec*G265::Cm^R^ *rec*O*ssb*-113^(ts)^, Therefore, we concluded that formations of RecG-yfp protein foci in live *E.coli* cells required also an intact C-terminus of SSB [22, 33, 62].

### Increased expression of RecG enhanced the UV resistance only in the absence of *recO*

To further understand the biological significance of our observations of RecG foci in the *rec*O null mutant, we compared the UV resistances of JM83*rec*F::Kan^R^, JM83*rec*R::Kan^R^ and JM83*rec*O::Kan^R^ mutant carrying *rec*G-pUC18 plasmid. We expected to see differences in UV resistance between JM83*rec*O::Kan^R^ JM83*rec*F::Kan^R^ and JM83*rec*R::Kan^R^ when supplementing with RecG. The results show that the viability of JM83*rec*O::Kan^R^ mutants was increased by supplementing RecG using plasmids of *rec*G-pUC18 (increased by 1.65-fold, Figure4D) and *rec*G-yfp-pUC18 (Figure 4E). In contrast, the viability of either JM83*rec*F::Kan^R^ or JM83*rec*R::Kan^R^ was unaffected by the supplementation of RecG (Figure 4A-C). This suggested that although RecO, RecF and RecR are thought to work epistatically in the RecF pathway of homologous recombination by forming either RecOR or RecFOR complexes [52], but a distinct role of RecO, RecR and RecF can be distinguished upon their binding with SSB[41, 60,61,62, 74]. Our observations that RecG-yfp foci required a functional SSB C-terminus only in the *rec*O::Kan^R^ cells rather than in the *rec*R::Kan^R^ cells or the *rec*F::Kan^R^ cells fits this paradigm. Although RecG participates in each of the three pathways of homologous recombination [18,19], some have suggested that RecG functions predominantly in the RecF pathway. For example, inactivation of *rec*G gene stimulated RecF pathway of homologous recombination instead of RecBCD pathway of homologous recombination, suggesting that RecG somehow inhibits the RecF pathway of homologous recombination either by competing with RecO binding to SSB, as mentioned, or by offering an alternative route to bypass the RecF pathway of homologous recombination repair [75,76].

**Figure 4.**
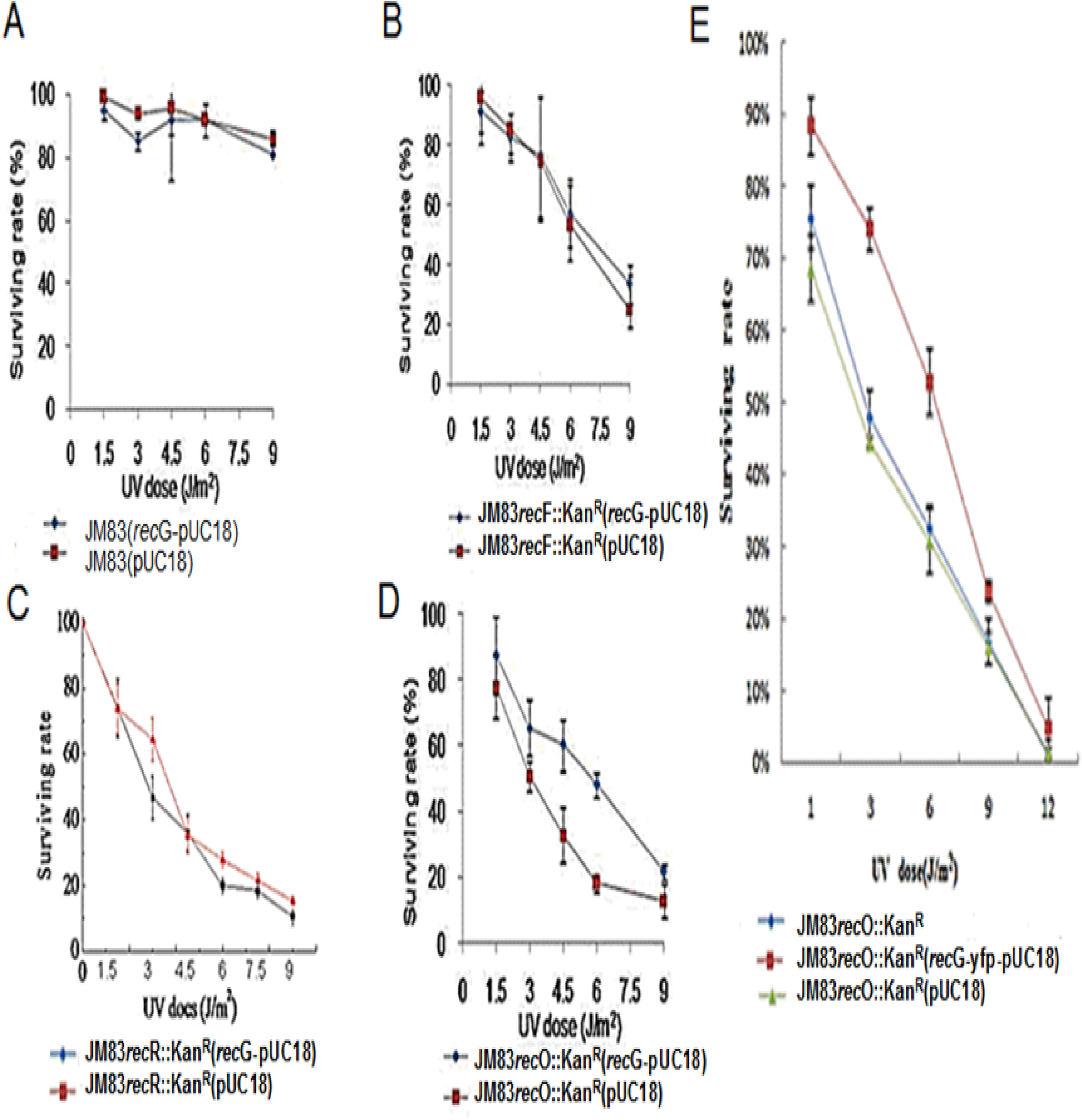
Significant increase in the viability of JM83*rec*O::Kan^R^ by expression of *rec*G. 4A, viability of JM83-WT when supplementing with RecG 4B, viability of JM83*rec*F::Kan^R^ when supplmenting with RecG 4C, viability of JM83*rec*O::Kan^R^ when supplmenting with RecG 4D, viability of JM83*recR*::Kan^R^ when supplementing with RecG 4E, viability of JM83*rec*A::Cm^R^*recO*::*Kan*^*R*^ when supplementing with RecG

### RecG–dependent and RecO-dependent repair of DNA double strand breaks in an *ssb-113* mutant

To further understand the RecO blocking RecG in binding to the C-terminus of SSB *in vivo*, and the partial increase in UV resistance by supplementing RecG in the JM83*rec*O::Kan^R^ mutant, we have compared the UV resistance with two groups of strains including JM83, JM83*rec*G::Cm^R^, JM83*rec*O::Kan^R^, JM83*rec*G::Cm^R^*rec*O::Kan^R^, and JM83*ssb-113*, JM83*ssb-113rec*G::Cm^R^, JM83*ssb-113rec*O::Kan^R^, JM83*ssb-113rec*G::Cm^R^*rec*O::Kan^R^. Wang and Smith reported that introducing an *ssb-*113 in *E.coli* raised a significant inhibition on DNA synthesis and generation of double-strand breaks due to loss of protection of single-stranded parental DNA opposite daughter strand gaps from nuclease attack by SSB[67]. The results of such comparisons were shown in Figure 5.

**Figure 5.**
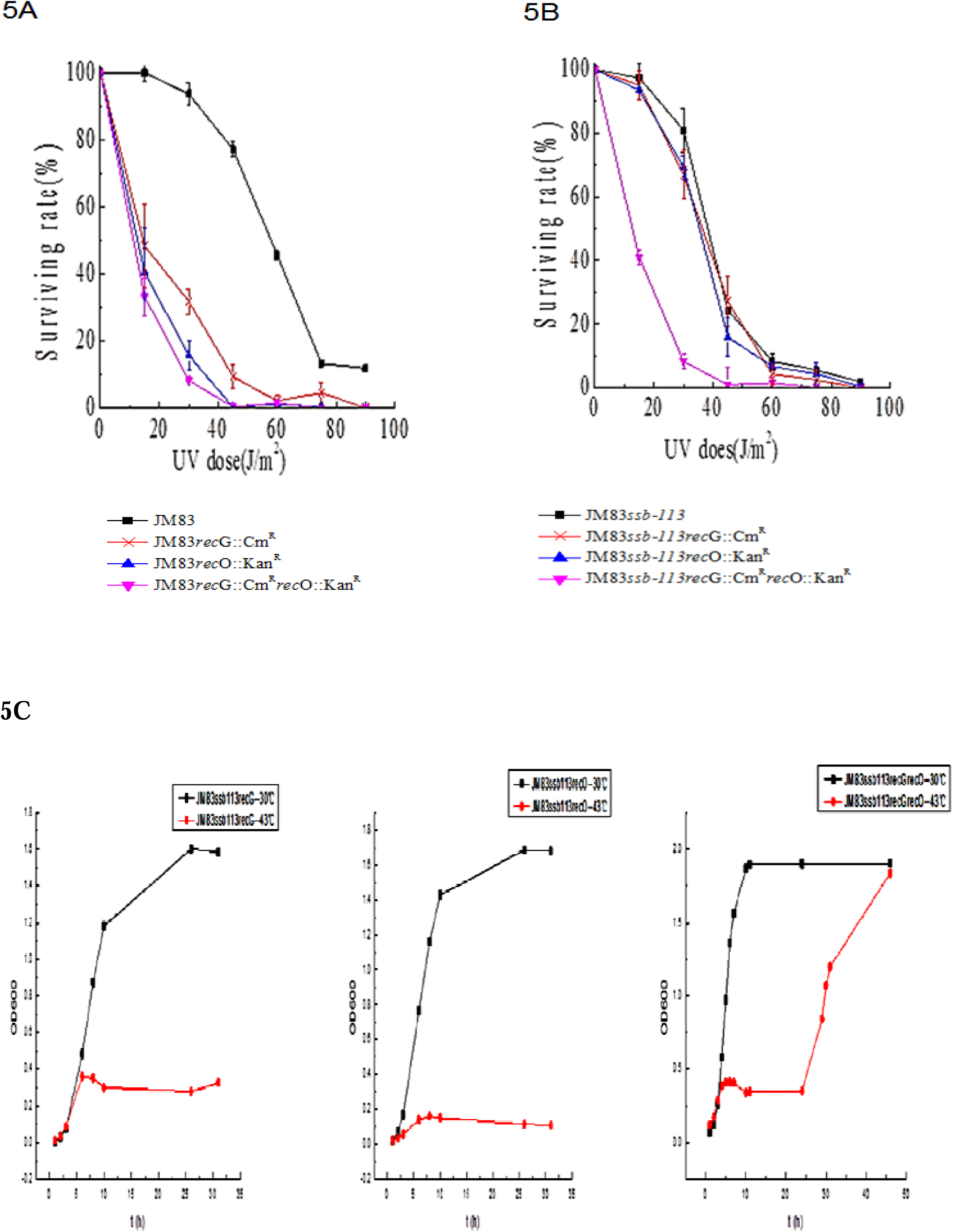
UV damage repair in different defection strains. 5A, *ssb* wildtype 5B, *ssb-113* mutant 5C, temperature-dependent growths of *ssb*-113 mutants defective with *rec*G and *rec*O,

The viability of strains carrying single *rec*G and *rec*O null mutations were similar to those of the strains carrying *rec*G and *rec*O double null mutations when a wildtype *ssb* gene was present in the cells. In contrast, the viability of the *ssb-113* strains carrying single *rec*G or *rec*O null mutations differed from those of the *ssb-113* strains carrying *rec*G and *rec*O double null mutations (Figure 5A and B). Strains carrying *recGssb-113* and *recOssb-113* double mutant genes showed similar UV resistance to that of the *ssb-113* single mutant when grown at 30°C after UV irradiations. However, the triple mutants of *rec*G, *rec*O and *ssb-113* showed an increased UV sensitivity than those of *ssb-113, recGssb-113* and *recOssb-113* mutant strains (Figure 5B), showing that RecG and RecO took part in the DNA replication in *ssb*-113 mutant cells.

In addition, we also compared the temperature dependent growth of JM83*ssb-113rec*G::Cm^R^, JM83*ssb-113rec*O::Kan^R^, and JM83*ssb-113rec*G::Cm^R^*rec*O::Kan^R^ at 30□ and 42□.. The results show that simultaneous inactivation of *rec*G and *rec*O improved the growth of JM83*ssb-113(ts)* at 42□ (Figure 5C). These observations put together suggested that RecG and RecO take part in the DNA double strand break repair in *ssb-113* mutant cells when grown at lower temperature (30□), while also generating DNA double strand breaks when grown at higher temperature (42□) (Figure 5C).

## Discussion

In the last decades, certain proteins/enzymes working in DNA repair and homologous recombination in *E.coli* have been suggested to target DNA replication forks via polymerase clamp or single-stranded DNA binding proteins [3, 22,49,50,53,63 and reference herein). Some of them have even been demonstrated to do so *in vitro*. For example, both RecG and RecO have been demonstrated to interact with the C-terminus of SSB *in vitro*, forming either RecG-SSB or RecO-SSB protein complex [17, 30,77]. However, visualizations of their interactions with a DNA replication fork in live *E.coli* cells have never been easy because of some reasons [64,69].

To understand how RecG and RecO may work in the interface of DNA replication, repair and recombination in *E.coli*, we have analyzed the protein foci of RecG-yfp in the *E.coli* cells absence of RecO, RecR, RecF, and also the wildtype SSB and SSB-113 mutant. We found that RecG-yfp formed fluorescent foci only in the absence of RecO when the C-terminus of SSB was intact, suggesting that RecO may compete with RecG in binding to the C-terminus of SSB in wildtype *E.coli* cells. To further understand the significance of this competition between RecG and RecO in the presence of wildtype SSB, we have carried out genetic analysis on the effects of RecG and RecO on the UV-resistance in an *ssb-113* mutant. The *ssb-113* gene used in this work is one of the two best-characterized *ssb* gene mutants that show temperature-dependent growth, another is *ssb-1*. The *E.coli* cells carrying *ssb-113* mutant gene were difficult to grow at 42^°^ C due to increased frequency on generating DNA double strand breaks by nucleases attacking the single stranded parental DNA template opposite to the newly synthesized daughter strand in a DNA replication fork, leading to profound UV-and ionizing radiation (IR) sensitivity even they were grown at permissive temperature [78]. Similar to SSB-deltaC8 mutant protein, SSB-113 protein showed very weak to no RecO binding capacity *in vitro* [41, 59, 63, 77, 79], and incapable of loading RecOR to facilitate the formation of ssDNA-RecA, as seen during homologous recombination, SOS response and DNA PolV catalyzed DNA translesion synthesis [22,80, 76].

By using a JM83*ssb*-*113* mutant, we found the UV resistance and the temperature dependent growth associated with proliferation of the JM83*ssb*-*113* mutant remain unaffected by inactivation of either *rec*G or *rec*O. Both JM83Δ*rec*G::Cam^R^*ssb-113* and JM83Δ*rec*O::Kan^R^*ssb*-113 mutants showed similar UV resistance and temperature dependent growth to those of an *ssb-113* alone mutant. However, simultaneous inactivation of RecG and RecO decreased the UV resistance of a JM83Δ*rec*GΔ*rec*O*ssb*-113 triple mutant when grown at 30□, and improved the temperature dependent growth of the JM83Δ*rec*G::Cam^R^Δ*rec*O::Kan^R^ *ssb*-113 mutant at 42□ (Figure 5). Our these observations argued that RecG and RecO worked in avoiding DNA double strand breaks generation in the JM83*ssb-113* cells when they were experiencing UV irradiation and grown at permissive temperature, while they might also work to make DNA double strand breaks when grown at nonpermissive temperature (Figure 6).

**Figure 6.**
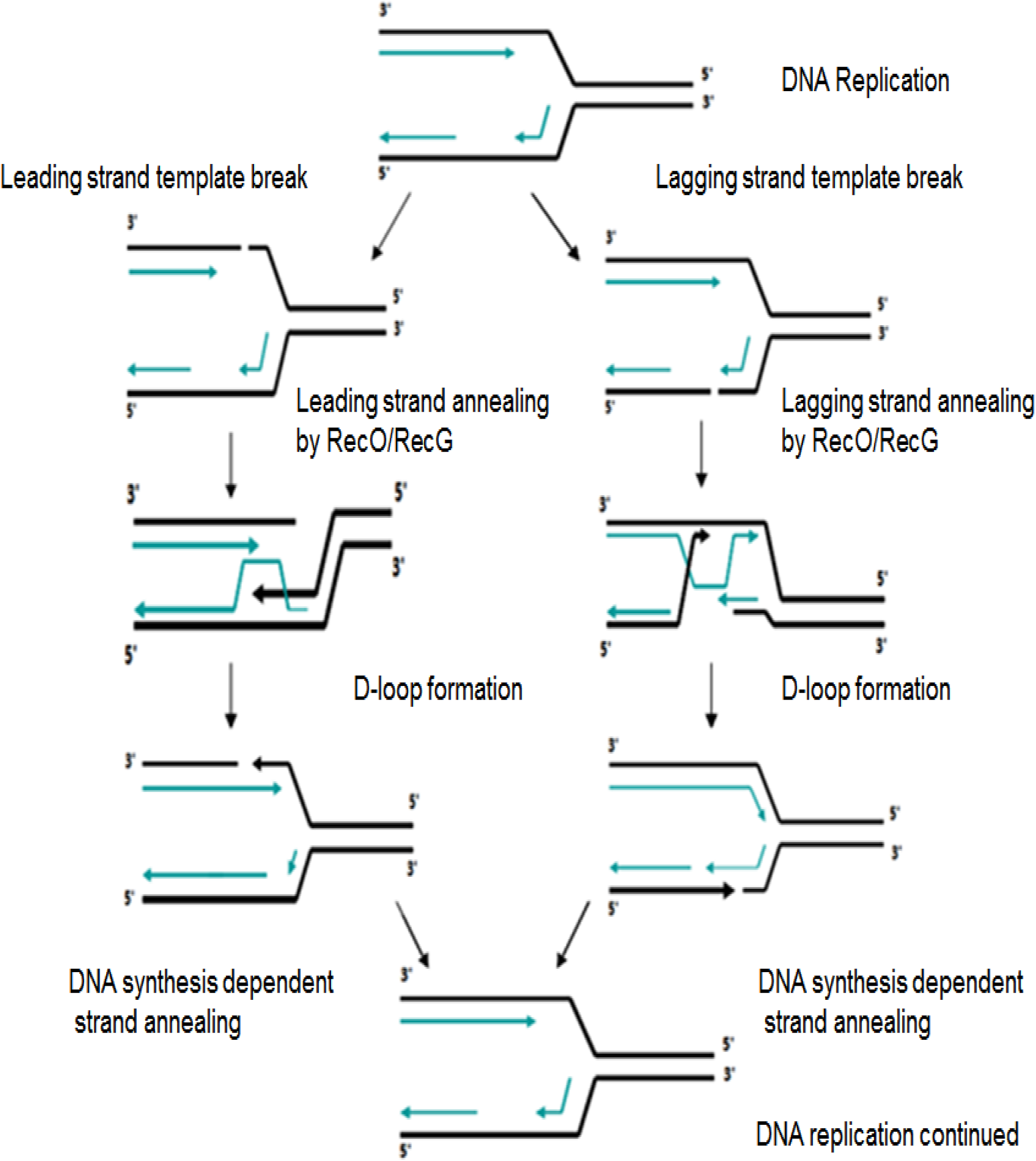
Schematic illustration for a role of RecO and RecG in DNA double strand break repair in a RecA-independent DNA Replication. RecG unwinds and rewinds the newly synthesized strands on the leading and lagging strand template of the DNA replication fork, alternatively, RecO anneals the SSB-113/SSB coated template strands into double strands. Each of these manipulations can work dependently and independently in a RecA-independent repair on DNA double strand breaks

Both RecG and RecO could participate in repair DNA double strand break through using DNA strand annealing/rewinding activities in a RecA-independent manner (Figure 6). RecO might be used to anneal the two parental template strands in a DNA replication fork by pairing two nascent strands in a problematic DNA replication fork [22, 34]; Alternatively, this might also be done by RecG unwinding and rewinding (Figure 6).

The proposed DNA strand annealing can be seen in a RecG-catalyzed fork regression event, or a DNA synthesis dependent strand annealing pathway of homologous recombination (SDSA). It has been well studied that the SDSA and the DSBR are two primary models of homologous recombination explaining how a DNA double stranded break could be repaired using homologous recombination[3, 47]. They resemble to each other in steps of DNA ends resection and DNA synthesis, but differ in how a 3’ overhang (which was not involved in strand invasion) formed a Holliday junction. In a conventional DSBR pathway, the 3’ overhang was suggested to form a Holliday Junction, while in an SDSA pathway, the 3’ overhang was suggested to be released from a D loop intermediate after it was extended using DNA polymerase, and which then to re-anneal with its complementary DNA segment (as depicted in Figure 6), leading to a small flap of DNA to be removed (Figure 6). The SDSA is frequently used to avoid crossover formation in meiotic recombination, which is believed to manage most non-crossover recombinants generated in eukaryotic meiosis, such as in *S. cerevisiae*. The conventional DSBR akin to that of *E.coli* turned to be the minor pathway in the eukaryotic homologous recombination repair of DNA double strand breaks, and presumably to be used only in mitotically proliferating cells [81]. This turned out to be that although all RMPs and recombinase are well conserved throughout bacteriophage to eukaryotes, they may work differently in the regulating a specific pathway of repair or homologous recombination [82]. Indeed, many lines of evidence showed that the strand annealing activity of RecO needed to be regulated by binding to single strand binding protein in divergent bacterium species, including *Deinococcus radiodurans, Mycobacteria, Bacillus subtilis* and *E.coli* [35, 63, 83, 84, 85]. For example, the RecO protein isolated from both *E.coli* and *Bacillus subtilis* was found to be able to bind SSB. Intriguingly, RecO proteins isolated from *D. radiodurans* and *Mycobacteria* have relinquished the SSB-binding, suggesting alternative linkages of DNA replication; repair and recombination existed in the interface of DNA replication, repair and homologous recombination [35,85].

In conclusion, our findings may be implicated with a mechanism of setting up a defined pathway of DNA repair and homologous recombination. More specifically, our studies may help understand why SDSA pathway of homologous recombination could be defined in organisms such as *Deinococcus radiodurans, Mycobacterium* and *S. cerevisiae* etc.[86, 87], but a similar SDSA pathway of homologous recombination has not been defined in *E.coli* until now.

## Materials and methods

### Chemicals and media

4’-6-Diamidino-2-phenylindole (DAPI) was purchased from Sigma Co., Ltd (Merck KGaA, Darmstadt, Germany). Restriction enzymes, plasmid extract kits and T4 DNA ligase were the products of Fermentas(Thermo Fisher Scientific, Waltham, MA USA). DNA recovery kits were purchased from Biomed (Los Angeles Biomedical Research Institute, USA). Isopropyl-beta-D-thiogalactopyranoside (IPTG) and 5-bromo-4-chloro-3-indolyl-beta-D-galacto-pyranoside (X-gal) were from Sino-American Biotechnology Co., Ltd. (Guang Dong, China). All PCR amplification primers used in this work were synthesized by Huada Gene Synthesis (Shenzhen, China). Luria-Bertani (LB) broth was purchased from Oxoid Ltd, (Hampshire, England). For cultivating the bacteria on plate, the LB broth was supplemented with 1.5% agar. All cultivations of the cells were carried out at 37°C except for those *ssb-113*(ts) mutants, which were cultivated at 30°C.

### Bacteria, phage and plasmids

*E.coli* strains used in this work were JM83 *(D Leach, Edinburgh*, UK) [88] and its derivatives. JJC451 *rec*F400::Tn5 (*Kan*^*R*^), JJC2135 *rec*O1504::Tn5 (*Kan*^*R*^), JJC2142 *rec*R252::Tn10 (*Kan*^*R*^),Δ*rec*G265: Tn10 (*Cm*^*R*^) (from B.Michel and M.Bichara in CNRS, France, respectively), and PAM2611 *Hfr (PO101), &lambda-, hisA323* (Stable), *ssb-113*(ts) (from CGSC, purchased by Yanbin Zhang, University of Miami). Strains of JM83*rec*F::*KanR*Δ*rec*G265::*Cm*^*R*^ (XP069), JM83*rec*O::*KanR*Δ*rec*G265::*Cm*^*R*^ (XP070), JM83*rec*R::*KanR*Δ*rec*G265::*Cm*^*R*^ (XP071), JM83*rec*F400::Tn5 (Kan^R^), JM83*rec*O1504::Tn5(Kan^R^), JM83 *rec*R252::Tn10(Kan^R^), JM83*rec*O::*Kan*^R^Δ*rec*G265::*Cm*^*R*^ *ssb*-113^(ts)^, JM83*rec*G::Cm^R^, JM83*rec*O::Kan^R^, JM83*rec*O::*Kan*^R^as::Tn10, JM83*rec*G::Cm^R^ *recO*::*Kan*^R^, JM83*rec*A::Cm^R^ *recO*::*Kan*^R^,JM83*rec*A::Cm^R^ *recR*::*Kan*^R^, JM83*rec*A::Cm^R^ *rec*F::Kan^R^, JM83*ssb-113*, JM83*ssb-113rec*G::Cm^R.^, JM83*ssb-113rec*O:: Kan^R^, and JM83*ssb-113rec*G::Cm^R^*rec*O::Kan^R^ etc were isogenic derivatives of JM83 constructed using P1 transduction[89]. Plasmids pUC18, *RecR-yfp*-pUC18, *MutS-gfp*-pACYC184, and pUC18-*yfp* were stocks of this laboratory. P1 Phage was a gift from D Leach (University of Edinburgh).

### Constructions of plasmids recG-yfp-pUC18 and recG-pUC18

Expression vectors of *RecG-yfp*-pUC18, *rec*G-pUC18 were constructed as follows: (1) for the construction of *RecG-yfp*-pUC18, PCR amplification of *rec*G gene of JM83 genome was conducted by using DNA primers: RecG_F: GCAGG<u>CTGCAG</u>GGTAAGTGC (underlined sequence is a *Pst* I restriction site) and RecG_R: CCGC<u>GGATCC</u>CGCATTCGAGTAACG (underlined sequence is a *Bam*H I restriction site). Fusion of *rec*G and *yfp* genes was carried out by merging the PCR product of *rec*G gene with an *yfp* gene at the *Bam*H I restriction site (*yfp* gene has a *Bam*H I site at its N-terminus). Plasmid *RecG-yfp*-pUC18 was subsequently constructed by inserting the *RecG-yfp* fusion gene into the *Pst* I and *Hin*d III restriction sites of plasmid pUC18, of which a stop codon “TTA” was embedded in *Hin*d III restriction site); (2) for the construction of RecG expression vector *rec*G-pUC18, PCR amplification of the *rec*G gene was conducted by using DNA primers: RecG_F: GCAGG<u>CTGCAG</u>GGTAAGTGC (underlined sequence is a cutting site of *Pst* I) and RecG_R: CTGCCG<u>AAGCTT</u>ACGCATTCGAG (underlined sequence is a cutting site of *Hin*d III, which contains a stop codon, TTA). The *rec*G-pUC18 was then constructed by inserting the *rec*G at the *Pst* I and *Hin*d III sites of pUC18, respectively. The α-complementation was carried out during cloning manipulations on LB plates containing the corresponding antibiotics, 40mM IPTG and 20mM X-gal. The *rec*G-pUC18 and *RecG-yfp*-pUC18 were confirmed by DNA sequencing using an oligonucleotide: 5’-ATCCACATTGCCCTCCATC and by restriction digestion using *Pst* I, *Bam*HI and *Hin*d III, respectively. All manipulations were conducted by following Molecular Cloning [90].

### Transformation and purification of plasmids

Plasmids transformed different *E.coli* cells, including wild-type and its isogenic mutants using a CaCl_2_ method. Recoveries of plasmids from the transformants were performed by using a plasmid mini preparation kit. Confirmations for the plasmids were carried out by using restriction digestions and followed by resolution of the digested products on agarose gel (Sambrook and Manniatis, 1989).

### Functional analysis of RecG and RecG-yfp in Δ*recG265::Cm*^*R*^ mutants

Activities of RecG and RecG-yfp proteins expressed by *rec*G-pUC18 and *recG-yfp*-pUC18 were evaluated by their improvements on the UV resistances of Δ*rec*G265::Cm^R^ mutant, respectively. In brief, *recG-yfp*-pUC18 or *rec*G-pUC18 transformed Δ*rec*G265::Cm^R^ mutants, and cells carrying either *recG-yfp*-pUC18 or *rec*G-pUC18 were irradiated using UV light, and the viabilities of the cells were scored as follows: Petri dishes containing different strains were irradiated under a 20W ultraviolet lamp by a distance of 70 cm for different time. Then the irradiated strains were cultivated at 37°C in dark until visible colonies formed. The numbers of colonies grown in the irradiated plates and unirradiated plates were counted for obtaining the surviving rate [70].

### Visualizations of the fluoresce tagged fusion proteins *in vivo*

Visualizations of RecG-yfp in JM83, the wild-type and its isogenic derivatives were carried out by using a Leica laser scanning confocal microscopy (Leica).

### Measurements of UV resistances and comparisons of growth

UV resistances of different JM83 mutants were examined by using the method as described by *Qiu* and *Pan* [70] and the aforementioned. The comparisons of temperature dependent growth of JM83*ssb-113rec*G::Cm^R^, JM83*ssb-113rec*O::Kan^R^, and JM83*ssb-113rec*G::Cm^R^*rec*O::Kan^R^ at 30°C and 42°C was carried out by following the methods described by Miller [89].

## Acknowledgements

This work was supported by grants from Natural Science Foundation of Beijing Institute of Technology (Grant No. 1060050320804) and Beijing Natural Science foundation (5132014) to XP. The funders had no role in study design, data collection and analysis, decision to publish, or preparation of the manuscript. We are very much grateful to David Leach (University of Edinburgh) for stains and P1 phage; B. Michel (CNRS, France) for AB1157recF/O/R mutants. Marc Bichara (Ecole Supérieure de Biotechnologie de Strasbourg, CNRS) for Δ*rec*G265::Cm^R^ mutant, Yanbin Zhang for *ssb-113*(ts) and Li-jun Bi (Institute of Biophysics, Chinese Academy of Sciences) for a *mut*S-*gfp* fusion gene; Thanks also go to Piero Bianco (the State University of New York at Buffalo) for thorough reading, editing of the manuscript and also valuable discussion on this work, and Shuang Han, Shan Jian for technical assistance. LY, XSY and XFC are graduate students in the laboratory.

## Author Contributions

XP designed experiments, supervised the experiments, analyzed data and wrote the manuscript; LY, N J, XC, BL, XY, YD performed the experiments and analyzed the data, and LD and FD helped in organization, participate discussions.

